# Distinct molecular etiologies of male and female hepatocellular carcinoma

**DOI:** 10.1101/507939

**Authors:** Heini M. Natri, Melissa A. Wilson, Kenneth Buetow

## Abstract

Sex-differences in cancer occurrence and mortality are evident across tumor types; men exhibit higher rates of incidence and often poorer responses to treatment. Targeted approaches to the treatment of tumors that account for these sex-differences require the characterization and understanding of the fundamental biological mechanisms that differentiate them. Hepatocellular Carcinoma (HCC) is the second leading cause of cancer death worldwide, with the incidence rapidly rising. HCC exhibits a male-bias in occurrence and mortality, but previous studies have failed to explore the sex-specific dysregulation of gene expression in HCC. Here, we characterize the sex-shared and sex-specific regulatory changes in HCC tumors in the TCGA LIHC cohort. By using a sex-specific differential expression analysis of tumor and tumor-adjacent samples, we uncovered etiologically relevant genes and pathways differentiating male and female HCC. While both sexes exhibited activation of pathways related to apoptosis and cell cycle, males and females differed in the activation of several signaling pathways, with females showing PPAR pathway enrichment while males showed PI3K, 305 PI3K/AKT, FGFR, EGFR, NGF, GF1R, Rap1, DAP12, and IL-2 signaling pathway enrichment. Using eQTL analyses, we discovered germline variants with differential effects on tumor gene expression between the sexes. 24.3% of the discovered eQTLs exhibit differential effects between the sexes, illustrating the substantial role of sex in modifying the effects of eQTLs in HCC. The genes that showed sex-specific dysregulation in tumors and those that harbored a sex-specific eQTL converge in clinically relevant pathways, suggesting that the molecular etiologies of male and female HCC are partially driven by differential genetic effects on gene expression. Overall, our results provide new insight into the role of inherited genetic regulation of transcription in modulating sex-differences in HCC etiology and provide a framework for future studies on sex-biased cancers.

## Background

Differences in cancer occurrence and mortality between sexes are evident across tumor types; males exhibit higher rates of cancer incidence and often poorer response to treatment, including some forms of chemotherapy and immunotherapy [1,2]. While differences in risk factors may explain some portion of the sex-bias, the bias remains after appropriate adjustment for these factors [3,4]. A recent study examining the mutational profiles of tumors from males and females across The Cancer Genome Atlas (TCGA) found sex-differences in mutational profiles, calling for the consideration of sex as a biological variable in studies on cancer occurrence, etiology, and treatment [5]. Despite these underlying molecular differences, sex is rarely considered in the development of cancer therapies.

Across tumor types analyzed, the largest sex-differences in autosomal mutational profiles were seen in liver hepatocellular carcinoma (HCC), indicating that male and female HCC are etiologically distinct [5]. Furthermore, HCC exhibits sex-bias in occurrence, with a male-to-female incidence ratio between 1.3:1 and 5.5:1 across populations [6,7]. The sexes also differ in the clinical manifestation of HCC, males exhibiting an earlier onset and more/larger nodules [8]. HCC is the second leading cause of cancer mortality worldwide, accounting for 8.2% of all cancer deaths [2], and the incidence in the US has doubled in the last 3 decades, attributable to increased rates of obesity [7], calling for the development of new interventions and targeted therapies.

Sex-specific gene regulation may partially underlie differences between the sexes in disease prevalence and severity [9,10]. Previous work observed extensive sex-biased signatures in gene expression in HCC and other sex-biased cancers [11]. However, this study focused solely on comparing male and female tumor samples, without consideration sex-differences in non-diseased and tumor-adjacent tissues. To understand cancer-specific processes, it is necessary to contrast the sex-differences in gene expression identified in HCC with those in non-tumor and tumor tissues. For the targeted treatment of tumors, it is necessary to understand whether sex differences in cancer reflect unique cancer-specific changes, or are reflective of healthy sex-differences that may underlie observed sex-bias in cancer occurrence and disease etiology.

In addition to sex differences in overall gene expression due to the wide effects of sex as a biological variable, genetic variants may alter gene expression in a sex-specific manner. A pan-cancer analysis of the TCGA dataset identified 128 germline variants altering gene expression levels (eQTLs) in HCC [12]. However, this study purposefully controlled for and removed the effect of sex and, to date, a sex-specific eQTL analysis in HCC has not been performed. Sex-stratified analyses can reveal sex-biased genetic effects on gene expression that may be obscured in a joint analysis of both sexes - e.g. cases where the regulatory variant has a zero or very small effect in one sex, or the eQTL exhibits an opposite effect direction in the two sexes [13]. eQTLs that are discovered in one sex but not in the whole sample analysis are likely to affect gene expression in a sex-dependent manner, and while a combined analysis of both sexes achieves a greater statistical power to detect sex-shared effects, it dilutes the signal of sex-dependent effects [14].

Targeted approaches to the treatment of male and female HCC require the characterization and understanding of the fundamental biological mechanisms that differentiate them. Here, we analyzed data from TCGA and The Genotype-Tissue Expression project (GTEx) to examine the sex-specific patterns of gene expression and regulation in HCC. Here, we have contrasted the sex-biased patterns of gene expression in HCC tumors with healthy and tumor-adjacent liver tissues, allowing us to detect sex-differences in gene expression shared between and specific to the different tissues. We show that male and female HCC exhibit differences in the dysregulation of genes and the germline genetic regulation of tumor gene expression. Importantly, these orthogonal approaches identify genes that converge in shared pathways, indicating sex-specific etiology in HCC. The results presented here have implications for the development of targeted therapies for male and female HCC.

## Methods

### Data

GTEx (release V6p) whole transcriptome (RNAseq) data (dbGaP accession #8834) were downloaded from dbGaP. TCGA LIHC Affymetrix Human Omni 6 array genotype data, whole exome sequencing (WES) and RNAseq data (dbGaP accession #11368) were downloaded from NCI Genomic Data Commons [15]. In total, RNAseq data from 91 male and 45 female GTEx donors, germline genotypes and tumor RNAseq data from 248 male and 119 female TCGA LIHC donors, as well as paired tumor and tumor-adjacent samples from 28 male and 22 female TCGA LIHC donors were utilized in this study. FASTQ read files were extracted from the TCGA LIHC WES BAM files using the *strip_reads*() function of *XYAlign* [16]. We used *FastQC* [17] to assess the WES and RNAseq FASTQ quality. Reads were trimmed using *TRIMMOMATIC IlluminaClip* [18], with the following parameters: seed mismatches 2, palindrome clip threshold 30, simple clip threshold 10, leading quality value 3, trailing quality value 3, sliding window size 4, minimum window quality 30 and minimum read length of 50.

### Read mapping and read count quantification

Sequence homology between the X and Y chromosomes may cause the mismapping of short sequencing reads derived from the sex chromosomes and affect downstream analyses [16]. To overcome this, reads were mapped to custom sex-specific reference genomes using *HISAT2* [19]. Female samples were mapped to the human reference genome GRCh38 with the Y-chromosome hard-masked. Male samples were mapped to the human reference genome with Y-chromosomal pseudoautosomal regions hard-masked. Gene-level counts from RNAseq were quantified using *Subread featureCounts* [20]. Reads overlapping multiple features (genes or RNA families with conserved secondary structures) were counted for each feature.

### Germline variant calling and filtering

BAM files were processed according to Broad Institute *GATK* (*Genome Analysis Toolkit*) best practices [21–23]: Read groups were added with *Picard Toolkit*’s *AddOrReplaceReadGroups* and optical duplicates marked with *Picard Toolkit*’s *MarkDuplicates* (v.2.18.1, http://broadinstitute.github.io/picard/). Base quality scores were recalibrated with *GATK* (v.4.0.3.0) *BaseRecalibrator.* Germline genotypes were called from whole blood Whole Exome Sequence samples from 248 male and 119 female HCC cases using the scatter-gather method with *GATK HaplotypeCaller* and *GenotypeGVCFs* [21]. Affymetrix 6.0 array genotypes were lifted to GRCh38 using the UCSC *LiftOver* tool [24] and converted to VCF. Filters were applied to retain variants with a minimum quality score > 30, minor allele frequency > 10%, minor allele count > 10, and no call rate < 10% across all samples.

### Clinical characteristics and cellular content of tumor samples

Confounding effects, e.g. differences in clinical and pathological characteristics or cell type composition of the sequenced samples, may contribute to the observed effect modification when utilizing stratified analyses. We examined the differences in the clinical characteristics between males and females in the TCGA LIHC cohort. We used a *t*-test to test for the equality of means in patient age and cell type proportions, and Fisher’s exact test to test to detect differences in risk factors and pathological classifications (Supplementary Tables S1 and S2).

### Filtering of gene expression data

FPKM (Fragments Per Kilobase of transcript per Million mapped reads) expression values for each gene were obtained using *EdgeR* [25]. Each expression dataset was filtered to retain genes with mean FPKM≥0.5 and read count of ≥6 in at least 10 samples across all samples under investigation. In the comparative analysis of differentially expressed genes (DEGs) between the tumor vs. tumor-adjacent samples in males, females, and both sexes, genes that reached the previously described expression thresholds in at least one tissue in at least one sex were retained. This assures that the DEGs detected in the sex-specific and combined analyses are not due to filtering.

### Differential expression analysis

For differential expression (DE) analysis, filtered, untransformed read count data were quantile normalized and logCPM transformed with *voom* [26]. From the TCGA LIHC dataset, paired tumor and tumor-adjacent samples were available for 22 females and 28 males. From the GTEx liver dataset, 91 male and 45 female samples were used in the DE analysis. A multi-factor design with sex and tissue type as predictor variables were used to fit the linear model. *duplicateCorrelation* function was used to calculate the correlation between measurements made between tumor and tumor-adjacent samples on the same subject, and this inter-subject correlation was accounted for in the linear modeling. As the paired tumor samples differed significantly between the sexes in terms of race, tumor grade, and HBV status, (Supplementary Tables S1 and S2), these parameters were included in the linear models as covariates. Due to missing values in the covariate data, the final numbers of sample pairs used in the analyses were 18 females and 26 males.

DEGs between comparisons were identified using the *limma*/*voom* pipeline [26] by computing empirical Bayes statistics with *eBayes*. An FDR-adjusted *p*-value threshold of 0.01 and an absolute log2 fold-change (FC) threshold of 2 were used to select significant DEGs.

To reliably detect genes that are expressed in a sex-biased way in HCC but not in non-diseased liver or in tumor-adjacent tissue, we examined genes that were DE in the male vs. female tumor comparison using the previously described significance thresholds, but not in the male vs. female comparisons of normal or tumor-adjacent samples with a relaxed significance threshold of FDR-adjusted *p*-value ≤ 0.1 and absolute log2(FC) ≥ 0.

To detect genes that are dysregulated in tumors compared to matched tumor-adjacent samples in each sex, we identified DEGs in the tumor vs. tumor-adjacent comparison of males, females, and in the whole sample. DEGs that were identified in one sex but not in the other or in the combined analysis of both sexes were considered sex-specific. DEGs identified in the combined analysis were considered sex-shared. This approach allows the identification of high-confidence sex-specific events that are a result of the underlying biological differences as opposed to sampling or statistical power. ANOVA and Kruskal-Wallis tests were used to test for equality of fold changes of sex-shared and sex-specific DEGs across male, female, and all samples.

### Overrepresentation of biological functions and canonical pathways

We further analyzed the sex-shared and sex-specific tumor vs. tumor-adjacent DEGs as well as the sex-specific eQTL target genes (eGenes) to identify sex-shared and sex-specific pathways driving HCC etiology. We used the *NetworkAnalyst* web tool [27], which utilizes a hypergeometric test to compute *p*-values for the overrepresentation of genes in regards to GO terms and KEGG and Reactome pathways. An FDR-adjusted *p*-value threshold of 0.01 was used to select significantly overrepresented GO terms and canonical pathways.

### Accounting for confounding effects and population structure

Gene expression values are affected by genetic, environmental, and technical factors, many of which may be unknown or unmeasured. Technical confounding factors introduce sources of variance that may greatly reduce the statistical power of association studies, and even cause false signals [28]. Thus, it is necessary to account for known and unknown technical confounders. This is often achieved by detecting a set of latent confounding factors with methods such as principal component analysis (PCA) or Probabilistic Estimation of Expression Residuals (PEER) [29]. These surrogate variables are then used as covariates in downstream analyses. We derived 10 PEER factors from the filtered tumor gene expression data and used the weights of these factors as covariates in the eQTL analysis. We used the R package *SNPRelate* [30] to perform PCA on the germline genotype data. We accounted for population structure by applying the first three genotype PCs as covariates in the eQTL analysis.

### eQTL analysis

We used eQTL analyses to detect germline genetic effects on tumor gene expression. Similar to the DE analysis, we utilized combined and sex-stratified analyses to detect sex-shared and sex-specific effects. Germline genotypes and tumor gene expression data from 248 male and 119 female donors in the TCGA LIHC cohort were used in the eQTL analysis. Filtered count data was normalized by fitting the FPKM values of each gene and sample to the quantiles of the normal distribution. To account for technical confounders and population structure, 10 *de novo* PEER factors and three genotype principal components were used as covariates. *Cis*-acting (proximal) eQTLs were detected by linear regression as implemented in *QTLtools* v.1.1 [31]. Variants within 1Mb of the gene under investigation were considered for testing. We used the permutation pass with 10,000 permutations to get adjusted *p*-values for associations between the gene expression levels and the top-variants in *cis*: first, permutations are used to derive a nominal *p*-value threshold per gene that reflects the number of independent tests per cis-window. Then, *QTLtools* uses a forward–backward stepwise regression to determine the best candidate variant per signal [31]. FDR-adjusted *p*-values were calculated to correct for multiple phenotypes tested, and an adjusted *p*-value threshold of 0.01 was used to select significant associations. To allow the comparison of effect sizes of sex-specific and sex-shared eQTLs across the sexes, effects of each variant located within the 1Mb *cis*-window were obtained using the *QTLtools* nominal pass.

Similarly to the tumor vs. tumor-adjacent DEGs, eQTLs that were detected in one sex but not in the other or in the combined analysis were considered sex-specific, while eQTLs detected in the combined analysis were considered sex-shared. ANOVA and Kruskal-Wallis tests were used to test for equality of effect sizes of sex-shared and sex-specific eQTLs across male, female, and all samples.

### Estimating statistical power in the eQTL analysis

We used the R package *powereQTL* [32] to estimate the effect of the sample size to the statistical power to detect eQTLs in the combined analysis of both sexes and in the sex-specific analyses (Fig. S2).

### Genomic annotations of eQTLs

We used the R package *Annotatr* to annotate the genomic locations of eQTLs [33]. Variant sites were annotated for promoters, 5’UTRs, exons, introns, 3’UTRs, CpGs (CpG islands, CpG shores, CpG shelves), and putative regulatory regions based on ChromHMM [34] annotations.

## Results

### Sex-specific patterns of gene expression in HCC

We identified sex-differences in gene expression in non-diseased liver (GTEx; 21 sex-biased genes with an FDR-adjusted *p*-value ≤ 0.01 and an absolute log2FC ≥ 2), tumor-adjacent tissue (TCGA LIHC; 21 genes), and HCC (TCGA LIHC; 53 genes) to characterize the shared and unique sex-differences that may drive the observed sex-biases in HCC occurrence and etiology (Fig. 1, Supplementary Tables S3-5). X-linked *XIST* and Y-linked genes were expressed in a sex-biased way across all tissues. While sex-biased gene expression in non-diseased and tumor-adjacent tissues may contribute to the sex-differences in cancer occurrence, sex-biased expression in tumors is suggestive of distinct molecular etiologies of male and female HCC. We identified 34 genes that show sex-differences in expression in HCC, but not in tumor-adjacent tissue or non-diseased liver, even with a relaxed significance threshold (Fig. 1A). Notably, Notch-regulating *DTX1* (Fig. 1B) and signal transducer *CD24* were downregulated in male HCC.

**Fig. 1.**
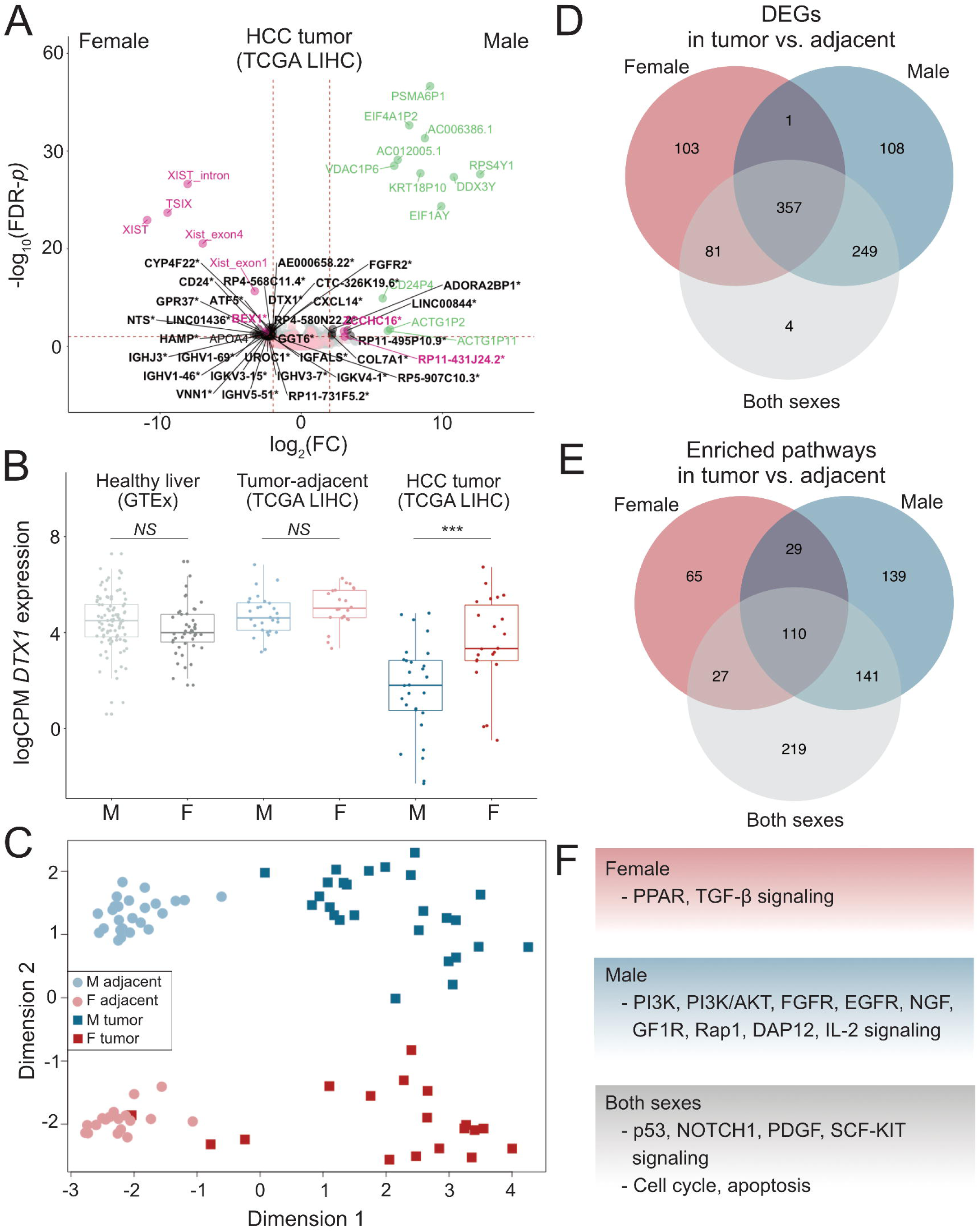
Patterns of gene expression and molecular etiologies of male and female HCC. **A:** Sex-biased gene expression in HCC. A volcano plot of DEGs between male (N=26) and female (N=18) HCC tumor samples. X-linked genes are indicated in pink, Y-linked in green, and autosomal in black. Significant genes were selected based on an FDR-adjusted *p*-value threshold of 0.01 and absolute log2(FC) threshold of 2. Multiple transcripts of the long non-coding RNA *XIST* are independently expressed. Genes that were not expressed in a sex-biased way in healthy liver (GTEx) or in the tumor-adjacent tissues are indicated with an asterisk. **B:** An example of a gene exhibiting a sex-bias in HCC but not in healthy liver or tumor-adjacent tissues. *DTX1* expression in log(CPM) is shown for male and female samples in each tissue. **C:** A multi-dimensional scaling plot of the paired TCGA LIHC tumor and tumor-adjacent samples of each sex. Euclidean distances between samples were calculated based on 100 genes with the largest standard deviations between samples. Tissue type (dimension 1) and sex (dimension 2) drive the overall patterns of gene expression in HCC. **D:** Venn-diagram of the overlap of DEGs in the sex-specific and combined analysis of matched tumor and tumor-adjacent samples. Substantially more DEGs were identified in the sex-specific analyses. **E:** Sex-specific and sex-shared DEGs were analyzed for the overrepresentation of functional pathways. Sex-specific patterns of pathway enrichment point to differential processes driving the etiology of male and female HCC. **F:** Examples of sex-specific and sex-shared pathways.

To further examine the sex-shared and sex-specific mechanisms driving HCC etiology, we detected DEGs between tumor and tumor-adjacent samples in males and females, as well as in the combined analysis of both sexes. Dimensionality reduction of gene expression data shows that variation among the tumor and tumor-adjacent samples is driven by tissue type and sex (Fig. 1C). When inspecting the tumor samples only, the first dimension is largely driven by sex (Supplementary Fig. S1). In the combined analysis of male and female samples, we detected 691 tumor vs. tumor-adjacent DEGs (Supplementary Table S6). In male- and female-specific analyses, we detected 715 and 542 tumor vs. tumor-adjacent DEGs, respectively (Supplementary Tables S7 and S8). Out of the total of 903 unique DEGs, 76.5% were shared between the sexes. We identified 103 female-specific and 108 male-specific tumor vs. tumor-adjacent DEGs. Notably, substantially more DEGs were detected in sex-specific analyses than in the unstratified analysis (Fig. 1D). Specifically, DEGs that showed different magnitudes in fold change between the sexes (based on ANOVA/Kruskal-Wallis tests) were detected in the sex-specific analyses (Fig. 2C, 2D), while DEGs with similar fold changes across all comparisons were detected in the combined analysis as well as the sex-specific analyses (Fig. 2A). Sex-shared DEGs that were only detected in the combined analysis, and not in the sex-specific analyses, showed a large variance in expression and, due to limited power, were not detected as statistically significant DEGs in sex-specific analyses (Fig. 2B). Tumor-infiltrating immune cells may produce spurious signals in DE analyses, which is evident from the detection of various immunoglobulin genes in tumor vs. tumor-adjacent comparisons (Supplementary Tables S6-8). However, male and female samples did not significantly differ in terms of cellular content (Supplementary Table S2), and thus such spurious signals are unlikely to affect male-female comparisons. The observed differences in gene expression are thus likely to reflect actual sex-differences rather than confounding differences in sample characteristics or composition.

**Fig 2.**
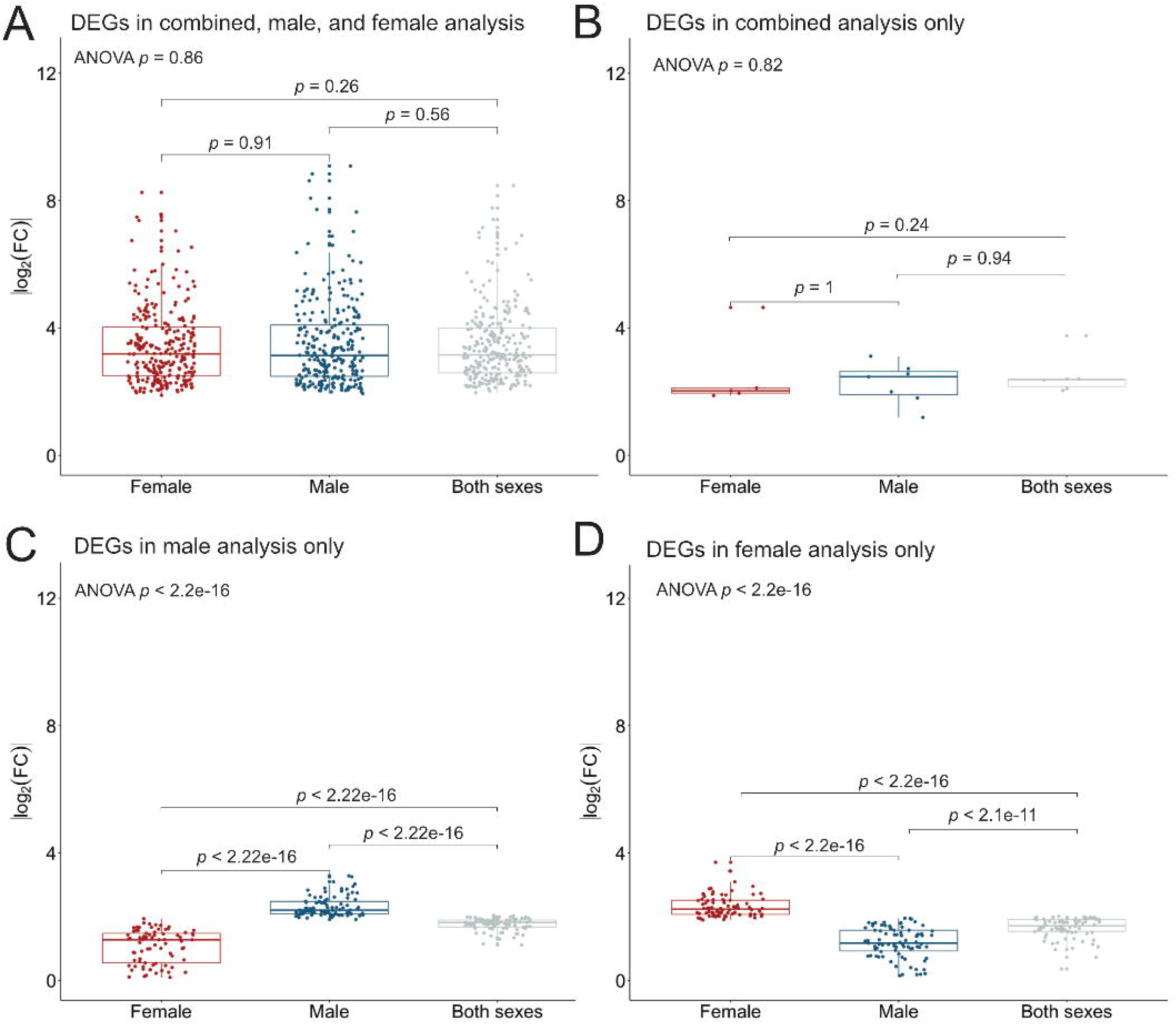
Absolute log2-fold changes of DEGs detected from tumor vs. tumor-adjacent comparisons in the sex-specific analyses and combined analysis of both sexes. Absolute log2-fold changes are given for female samples, male samples, and across all samples. Global *p*-values for ANOVA are shown for each DEG type. Adjusted *p*-values based on Kruskal-Wallis tests are shown for each pairwise comparison.

To put these results in a broader context, we analyzed the male- and female-specific DEGs (tumor vs. tumor-adjacent) for the overrepresentation of functional pathways. We found that the sex-shared and sex-specific DEGs were enriched in pathways relevant to oncogenesis and cancer progression (Supplementary Tables 9-11). We identified pathways that were overrepresented in only one of the sexes but not in the other or in the combined analysis of both sexes, indicating that male and female HCC are partially driven by different mechanisms and processes (Fig. 1E-F).

### Differential cis-eQTL effects in male and female HCC

To further investigate the mechanisms of sex-difference in HCC etiology, we used eQTL analyses to detect germline genetic effects on tumor gene expression in both the joint and sex-stratified analyses (Fig. 3A). We detected 1,204, 761, and 245 eQTLs in the combined, male-specific, and female-specific analyses, respectively (Supplementary Tables S12-14). As expected, genomic annotations show that most eQTLs are located on non-coding regions (Fig. 3B, Supplementary Tables S15-S17). Consistent with previous reports, most *cis*-eQTLs were located near transcription start sites (TSSs), with 63% of all eQTLs across the combined and sex-specific analyses being located within 20kb of TSSs. On average, 384 variants were tested per gene. 31% of the unique shared and sex-specific cis-eQTLs in HCC were also identified as eQTLs in the liver data in the GTEx project analysis release V7, indicating shared tissue origin. Out of the total of 1,595 unique associations, 75.7% were shared between the sexes. We detected 295 male-specific and 92 female-specific eQTLs. Since these associations were not detected in the unstratified analysis, they are likely not a result of differential power to detect associations due to different sample sizes, but exhibit effect modification by sex. Sex-specific associations exhibited differences in effect size between the sexes (based on ANOVA/Kruskal-Wallis tests, Fig. 4C, 4D), and the sex-specific effect is diluted in the combined analysis (Fig. 4C, 4D). Sex-shared large effect eQTLs were detected in sex-specific and combined analyses (Fig. 4A), and, due to the larger sample size, sex-shared low-effect eQTLs are detected in the combined analysis only (Fig. 4B).

**Fig. 3.**
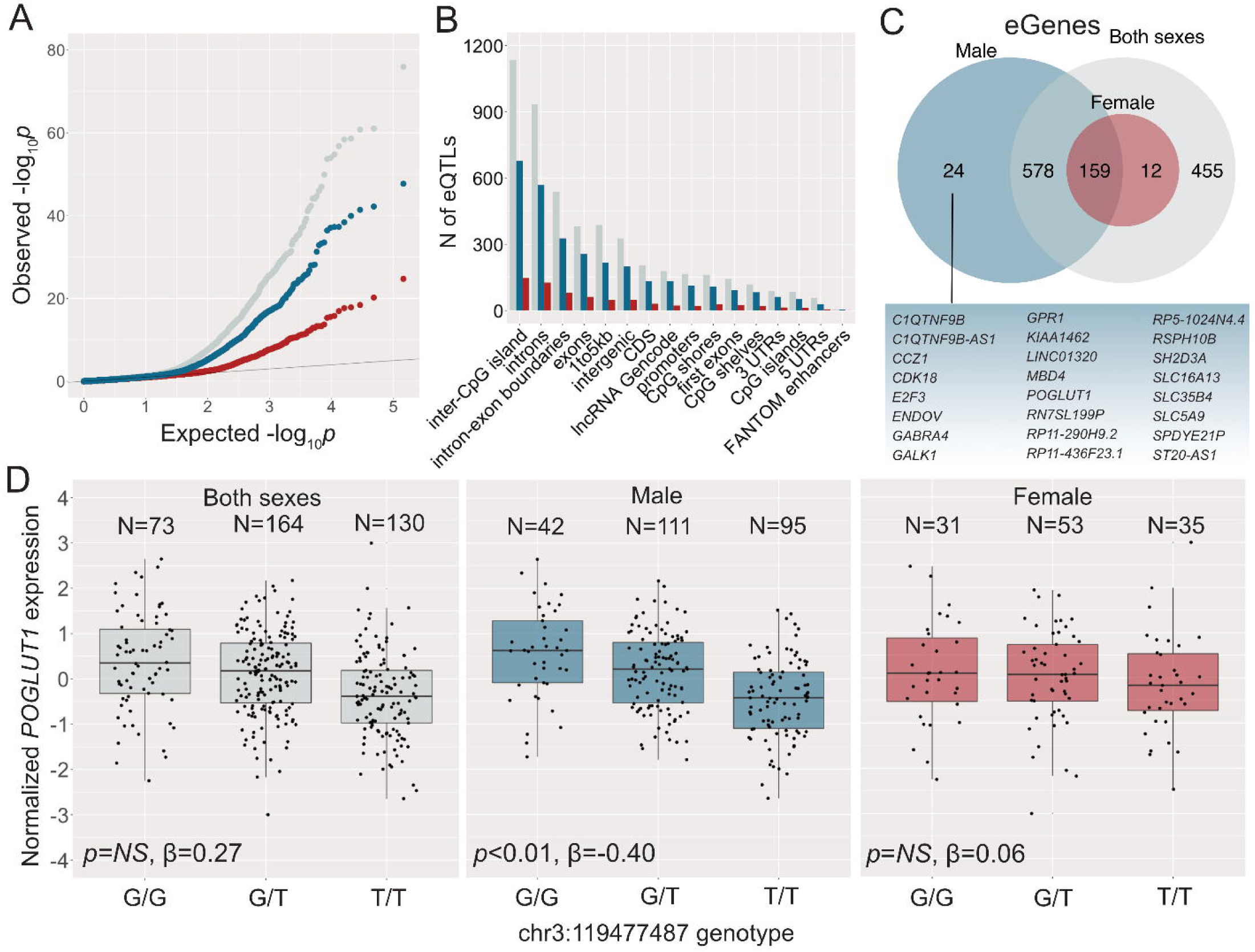
Sex-specific genetic effects on tumor gene expression in HCC. **A:** QQ-plot of eQTL associations in the combined analysis of both sexes (grey), male-specific analysis (blue), and female-specific analysis (red). **B:** Genomic annotations of eQTLs in the combined analysis of both sexes, male-specific analysis, and female-specific analysis. **C:** Overlap of eGenes detected in combined and sex-specific analyses. **D:** An example of a male-specific eQTL. *POGLUT1* expression in tumors is modulated by a germline variant in *cis* in male HCC, but not in female HCC nor in the combined analysis of both sexes, indicating effect modification by sex. Numbers of individuals with each genotype, adjusted significance, and effect size (β) are given for each model.

**Fig 4.**
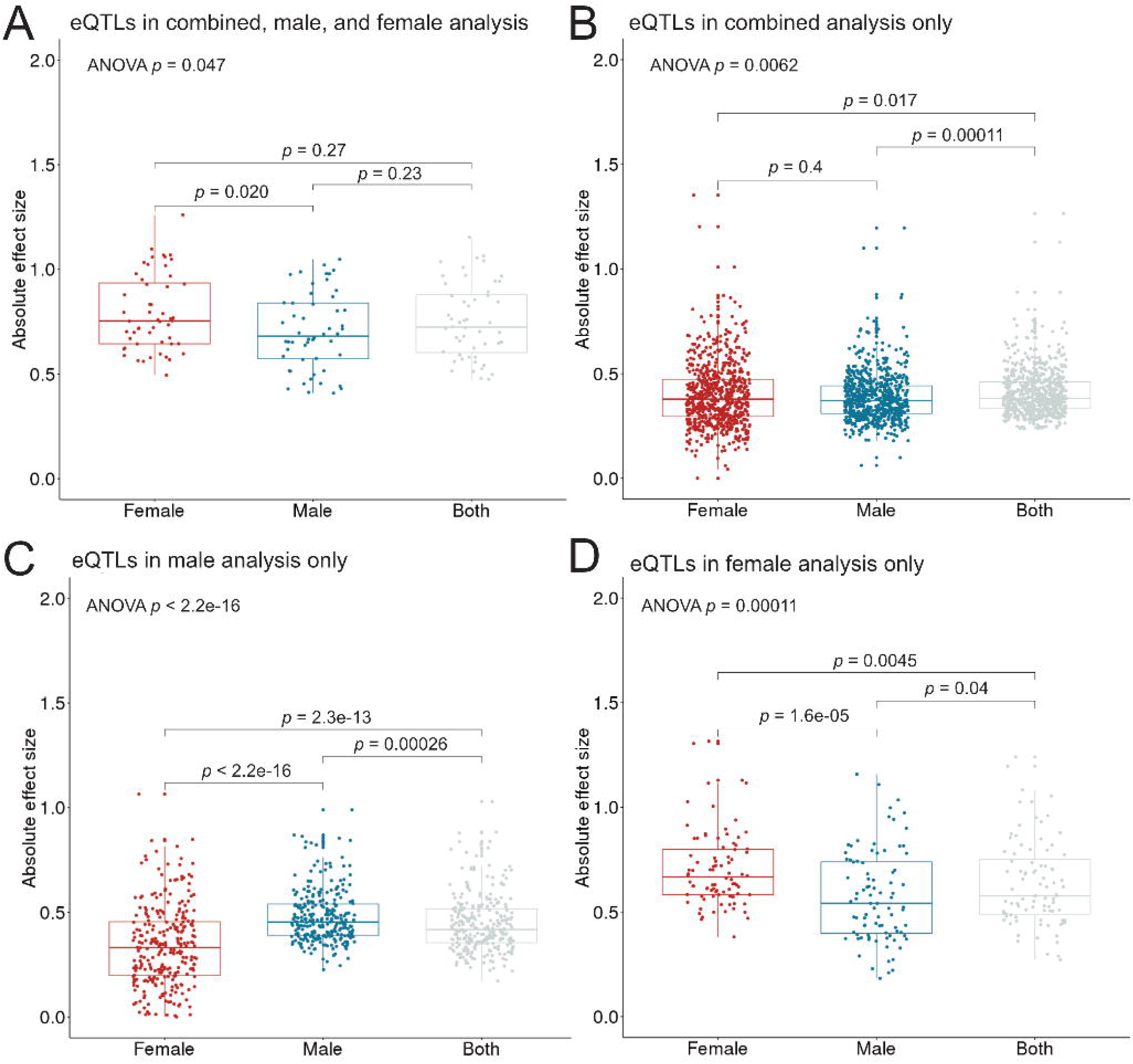
Absolute effect sizes of sex-shared and sex-specific eQTLs in males, females, and the whole study sample. Due to the larger sample size, sex-shared low-effect eQTLs are only detected as significant in the combined analysis (A). Sex-shared large effect eQTLs are detected in the combined analysis as well as the sex-specific analyses (B). Sex-specific eQTLs exhibit a larger effect in one sex than the other, and the effect is diluted in the combined analysis (C, D). Sex-shared large effect eQTLs can be detected in sex-specific and combined analyses. Global *p*-values for ANOVA are shown for each eQTL type. Adjusted *p*-values based on Kruskal-Wallis tests are shown for each pairwise comparison.

We detected 27 shared eGenes that were associated with independent variants in males and females. This could be due to actual biological differences in gene regulation, or due to technical constraints, in particular, missing genotypes in one sex affecting the permutation scheme to select the top-variant for each target gene. To overcome this and to detect high confidence instances of differential gene regulation between the sexes, we further examined the sex-shared and sex-specific eGenes: we found 24 genes that are under germline regulatory control in only male HCC (Fig. 3C), including *POGLUT1*, which is an essential regulator of Notch signaling (Fig. 3D). No genes were found to be associated with nearby variants in females only, likely due to reduced power to detect associations in females (Supplementary Fig. S2). Male-specific eGenes were overrepresented in pathways related to cell cycle, apoptosis, and cancer (Supplementary Table S18). Concordant with previous studies [14,35], none of the male-specific eGenes were differentially expressed between male and female HCC, indicating that the male-specific eQTLs are not a result of differences in overall gene expression levels between males and females, but are likely to arise from factors such as differential chromatin accessibility or transcription factor activity. The observation that non of the sex-biased autosomal genes in tumors harbor significant *cis*-eQTLs (Supplementary Table S19) also suggests that while sex-specific *cis*-eQTLs may contribute to differences in variance, sex-biased gene expression is likely a result of *trans*-effects, e.g. sex-chromosomal effects on autosomal gene expression, or, more widely, a result of sex as a biological variable, e.g. hormonal effects.

## Discussion

### Distinct molecular etiologies of male and female HCC

It is well established that patterns of gene expression vary between the sexes across different tissues. Previous studies have confounded these differences with those which may be driving etiological differences between male and female tumors. For example, Yuan et al. previously reported extensive sex-biased signatures in gene expression in HCC and other strongly sex-biased cancers [11]. While they identified immunity and cancer-associated enriched pathways based on sex-biased genes detected in HCC tumors, their approach was limited as it did not include the examination of non-diseased liver nor tumor-adjacent tissues. From the results presented here, we are able to distinguish the differences detected in comparisons of male and female HCC from those reflecting sex-differences in the healthy liver or in tumor-adjacent tissue, as well as to detect genes that are dysregulated in HCC in a sex-shared or sex-specific manner.

We characterized differences in gene expression between male and female HCC cases. Notably, sex-differences in gene expression were the largest in the tumor tissue, with 53 genes (including 32 autosomal genes) being expressed in a sex-biased way. These sex-differences point to distinct mechanisms underlying HCC oncogenesis between the sexes, and may partially underlie the observed sex-biases in HCC occurrence and onset. We detected 34 genes that were expressed in a sex-biased way in HCC tumors, but not in healthy or tumor-adjacent liver tissues. Some of these genes are of particular interest in the context of HCC: Notch-regulating *DTX1*, found here to be underexpressed in males compared to females, has been identified as a putative tumor suppressor gene in head and neck squamous cell carcinoma [36]. Another female-biased gene detected here, *CD24*, has a crucial role in T cell homeostasis and autoimmunity [37]. The opposing roles of *CD24* expression in cancer and autoimmune diseases raise interesting questions on the role of sex-differences in immunity underlying sex-differences in cancer. Future studies will focus on better understanding the differential regulation of immune functions between the sexes, and how these differences contribute to the observed biases in disease occurrence and etiology.

By sex-specific analyses of matched tumor and tumor-adjacent samples, we detected genes that are uniquely dysregulated in male and female HCC. Further examination of these genes revealed sex-differences in the pathway activation, indicating that the molecular etiologies of male and female HCC are partly driven by distinct functional pathways. Males and females differed in the activation of several signaling pathways, with females showing PPAR pathway enrichment while males showed PI3K, PI3K/AKT, FGFR, EGFR, NGF, GF1R, Rap1, DAP12, and IL-2 signaling pathway enrichment (Fig. 1E, Supplementary Tables 9-10). As these signaling pathways are notable targets for anti-cancer and anti-metastasis therapies [38–44], the results presented here have implications for the targeted treatment of male and female HCC.

### Sex-specific germline genetic effects on tumor gene expression may drive the molecular etiologies of male and female HCC

Sex-specific regulatory functions may underlie sex-differences in cancer etiology, progression, and outcome. We detected sex-differences in the germline genetic regulation of tumor gene expression in HCC, including 24 genes that were under germline regulatory control only in male HCC (Fig. 3). Functional annotations of these male-specific eGenes provide insight into possible regulatory mechanisms contributing to the observed male-bias in HCC and sex-differences in HCC etiology. Protein O-glucosyltransferase 1 (*POGLUT1*) was found to be under germline regulation in male HCC, but not in female HCC or in the joint analysis of both sexes (Fig. 3D). The eQTL associated with *POGLUT1* is located on a promoter region of its target (Supplementary Table S15). *POGLUT1* is an enzyme that is responsible for O-linked glycosylation of proteins. Altered glycosylation of proteins has been observed in many cancers [45,46], including liver cancer [47,48]. *POGLUT1* is an essential regulator of Notch signaling and is likely involved in cell fate and tissue formation during development. Genes involved in Notch and PI3K/AKT signaling were also found to be expressed in a sex-biased way in HCC tumors and overrepresented among the male-specific DEGs detected in the tumor vs. tumor-adjacent comparison, showing that sex-specific eQTLs and sex-specific dysregulated genes converge in canonical pathways. Notch signaling pathway was also detected as overrepresented (FDR-adj. *p*-value ≤ 0.01) among the 24 male-specific eGenes. PI3K-AKT is known to co-operate with Notch by triggering inflammation and immunosuppression [49]. These results point to a major role of the Notch/PI3K/AKT axis in the development of HCC in males. PI3K/AKT/mTOR signaling is of particular interest in the context of HCC, as it has been implicated in HCC carcinogenesis [50], is involved in hepatic gluconeogenesis [51], and is activated in a sex-biased way in the liver and other tissues [52]. The role of Notch and PI3K/AKT signaling in HCC may differ between early and late-stage tumors and among molecular subtypes, and further studies are necessary to understand the possible oncogenic properties of these pathways among HCC subtypes and between the sexes. In the future, analyses of data collected as a part of the International Cancer Genomics Consortium project may elucidate the sex-specific processes of HCC oncogenesis among the Japanese, as well as the interactions between sex and hepatitis infections in shaping HCC etiology. However, each dataset has a unique ancestry composition and are not directly comparable for validation purposes.

## Conclusions

In summary, we discovered differential regulatory functions in HCC tumors between the sexes. This work provides a framework for future studies on sex-biased cancers. Further studies are required to identify and validate sex-specific genetic effects on tumor gene expression and its consequences in HCC and other sex-biased cancers across diverse populations.

## Abbreviations

HCC: Hepatocellular Carcinoma.
TCGA: The Cancer Genome Atlas.
GTEx: Genotype x Tissue Expression Project.
HBV: Hepatitis B virus.
HCV: Hepatitis C virus.
DEG: Differentially expressed gene.
eQTL: Expression quantitative trait loci.
TSS: Transcription start site.

## Declarations

### Availability of data and materials

Data used in this study are available at dbGaP NCI Genomic Data Commons.

### Authors’ contributions

Conception and design: HMN, MAW, KB. Development of methodology: HMN, MAW. Acquisition of data: MAW, KB. Analysis and interpretation of data: HMN. Writing, review, and/or revision of the manuscript: HMN, MAW, KB. Study supervision: MAW, KB. All authors approved the final manuscript.

### Ethics approval and consent to participate

Not applicable.

### Consent for publication

Not applicable.

## Acknowledgments

We thank Dr. Nicholas Banovich and anonymous reviewers for their feedback on previous versions of this manuscript.

## Additional files

### Additional files contain supplementary information with two figures, and 19 supplementary tables

**Fig S1.**
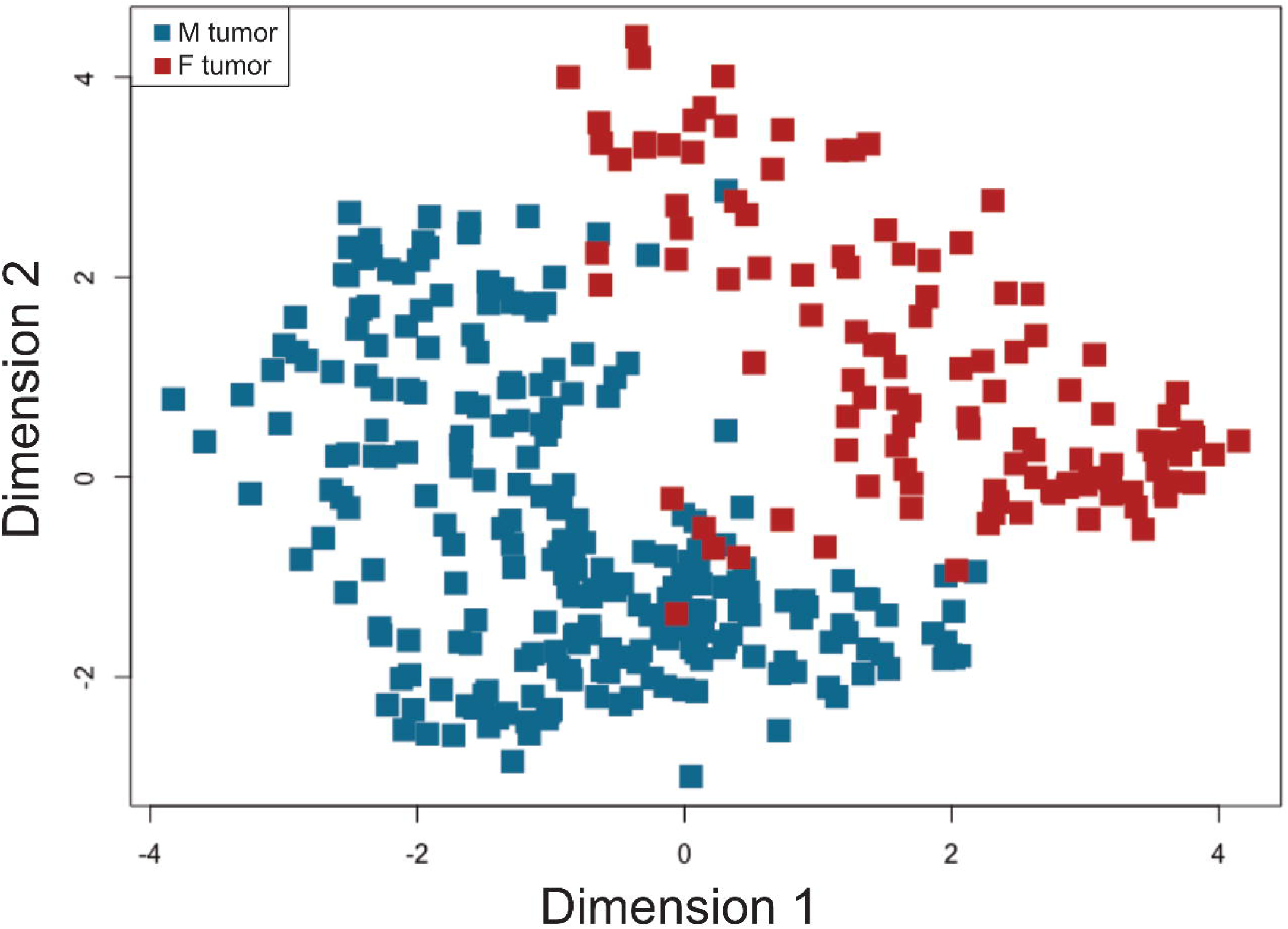
A multi-dimensional scaling plot of the TCGA LIHC tumor samples of each sex (N male = 248, N female = 119). Euclidean distances between samples were calculated based on 100 genes with the largest standard deviations between samples.

**Fig S2.**
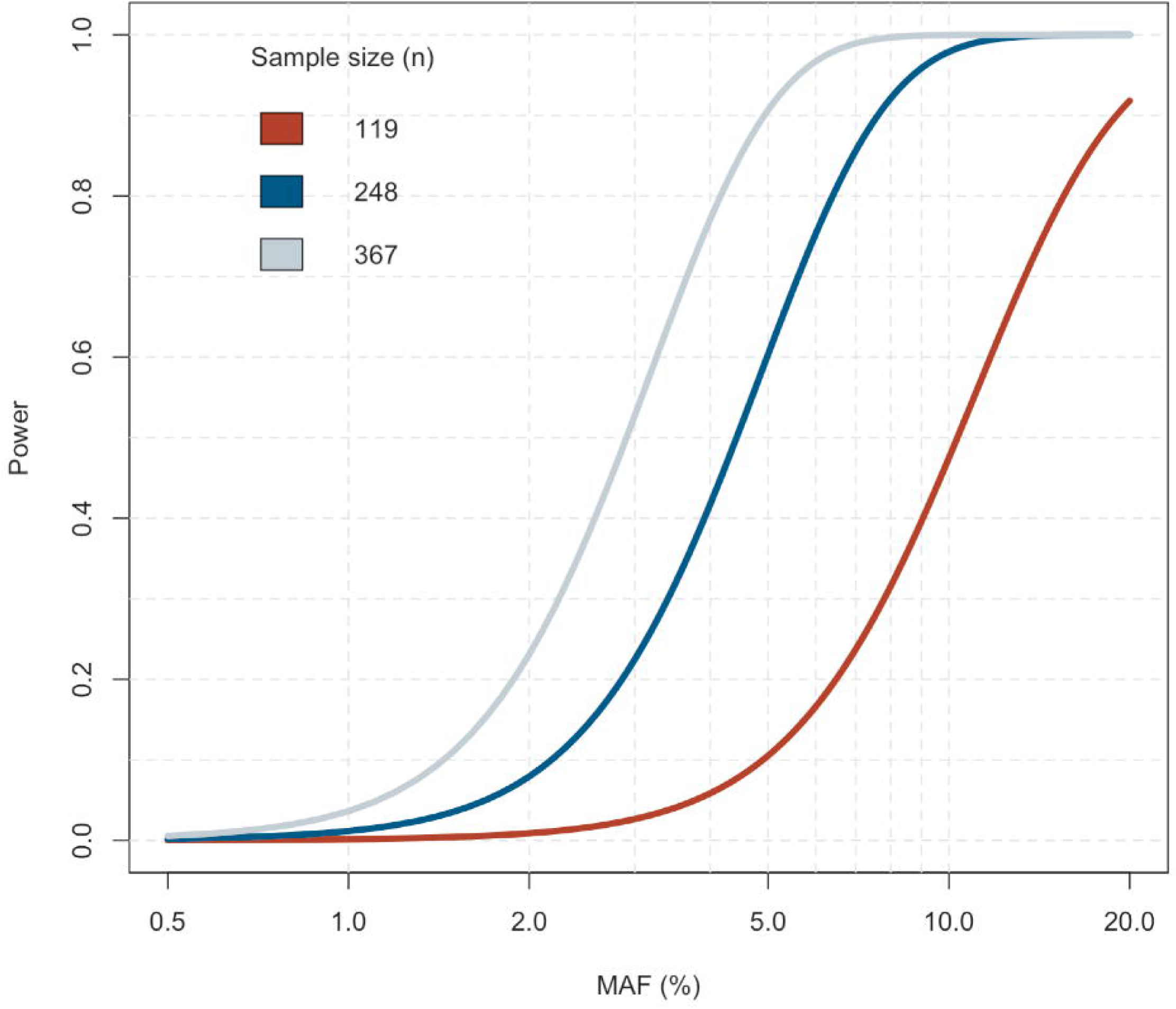
Estimation of statistical power in the combined (grey), male-specific (blue), and female-specific (red) eQTL analyses with a *p*-value level 0.01 and 384 variants. Increased power in the combined analysis allows the detection of sex-shared low-effect eQTLs.

**Table S1.** Categorical clinical characteristics of male and female patients in the TCGA LIHC cohort. Table S1. Categorical clinical characteristics of samples derived from male and female patients in the TCGA LIHC cohort. The following symbols are used to indicate statistical significance: * = p ≤ 0.10, ** = p ≤ 0.05, *** = p ≤ 0.01.

**Table S2.** Continuous clinical characteristics of male and female patients in the TCGA LIHC cohort.

**Table S3.** Sex-biased gene expression in the GTEx liver tissue samples with an FDR-adjusted *p*-value ≤ 0.1 and absolute log2 fold-change ≥ 0. N males = 91, N females = 45.

**Table S4.** Sex-biased gene expression in the TCGA LIHC tumor-adjacent samples with an FDR-adjusted *p*-value ≤ 0.1 and an absolute log2 fold-change ≥ 0. N males = 22, N females = 18.

**Table S5.** Sex-biased gene expression in TCGA LIHC tumor samples with an FDR-adjusted *p*-value ≤ 0.01 and an absolute log2 fold-change ≥ 2. N males = 22, N females = 18.

**Table S6.** Differentially expressed genes between matched tumor and tumor-adjacent samples in the combined analysis of male and female samples with an FDR-adjusted *p*-value ≤ 0.01 and an absolute log2 fold-change ≥ 2. N male sample pairs = 22, N female sample pairs = 18.

**Table S7.** Differentially expressed genes between matched male tumor and tumor-adjacent samples with an FDR-adjusted *p*-value ≤ 0.01 and an absolute log2 fold-change ≥ 2. N of sample pairs = 22.

**Table S8.** Differentially expressed genes between matched female tumor and tumor-adjacent samples with an FDR-adjusted *p*-value ≤ 0.01 and an absolute log2 fold-change ≥ 2. N of sample pairs = 18.

**Table S9.** Overrepresented GO terms and canonical pathways in the sex-shared tumor vs. tumor-adjacent DEGs. Significant terms and pathways were selected based on an FDR-adjusted *p*-value threshold of 0.01.

**Table S10.** Overrepresented GO terms and canonical pathways in the male-specific tumor vs. tumor-adjacent DEGs. Significant terms and pathways were selected based on an FDR-adjusted *p*-value threshold of 0.01.

**Table S11.** Overrepresented GO terms and canonical pathways in the female-specific tumor vs. tumor-adjacent DEGs. Significant terms and pathways were selected based on an FDR-adjusted *p*-value threshold of 0.01.

**Table S12.** *cis*-eQTLs detected in the combined analysis of both sexes. N=367 (N males = 248, N females = 119). Significant eQTLs were selected based on an FDR-adjusted *p*-value threshold 0.01.

**Table S13.** *cis*-eQTLs detected in the male-specific analysis. N=248. Significant eQTLs were selected based on an FDR-adjusted *p*-value threshold 0.01.

**Table S14.** *cis*-eQTLs detected in the female-specific analysis. N=119. Significant eQTLs were selected based on an FDR-adjusted *p*-value threshold 0.01.

**Table S15.** Genomic annotations of eQTLs detected in the combined analysis of both sexes.

**Table S16.** Genomic annotations of eQTLs detected in the male-specific analysis.

**Table S17.** Genomic annotations of eQTLs detected in the female-specific analysis.

**Table S18.** Overrepresented canonical pathways in the male-specific eQTL target genes. Significant terms and pathways were selected based on an FDR-adjusted *p*-value threshold of 0.01.

**Table S19.** Top-variants associated with autosomal genes that were expressed in a sex-biased way in HCC tumor samples.

## References

1. Clocchiatti A, Cora E, Zhang Y, Dotto GP. Sexual dimorphism in cancer. Nat Rev Cancer. 2016;16: 330.

2. Bray F, Ferlay J, Soerjomataram I, Siegel RL, Torre LA, Jemal A. Global cancer statistics 2018: GLOBOCAN estimates of incidence and mortality worldwide for 36 cancers in 185 countries. CA Cancer J Clin. 2018; doi:10.3322/caac.21492

3. Wisnivesky JP, Halm EA. Sex differences in lung cancer survival: do tumors behave differently in elderly women? J Clin Oncol. 2007;25: 1705–1712.

4. OuYang P-Y, Zhang L-N, Lan X-W, Xie C, Zhang W-W, Wang Q-X, et al. The significant survival advantage of female sex in nasopharyngeal carcinoma: a propensity-matched analysis. Br J Cancer. 2015;112: 1554–1561.

5. Li CH, Haider S, Shiah Y-J, Thai K, Boutros PC. Sex Differences in Cancer Driver Genes and Biomarkers. Cancer Res. 2018;78: 5527–5537.

6. Wands J. Hepatocellular carcinoma and sex. N Engl J Med. 2007;357: 1974–1976.

7. Petrick JL, Braunlin M, Laversanne M, Valery PC, Bray F, McGlynn KA. International trends in liver cancer incidence, overall and by histologic subtype, 1978-2007. Int J Cancer. 2016;139: 1534–1545.

8. Ladenheim MR, Kim NG, Nguyen P, Le A, Stefanick ML, Garcia G, et al. Sex differences in disease presentation, treatment and clinical outcomes of patients with hepatocellular carcinoma: a single-centre cohort study. BMJ Open Gastroenterol. 2016;3: e000107.

9. Gilks WP, Abbott JK, Morrow EH. Sex differences in disease genetics: evidence, evolution, and detection. Trends Genet. 2014;30: 453–463.

10. Morrow EH. The evolution of sex differences in disease. Biol Sex Differ. 2015;6: 5.

11. Yuan Y, Liu L, Chen H, Wang Y, Xu Y, Mao H, et al. Comprehensive Characterization of Molecular Differences in Cancer between Male and Female Patients. Cancer Cell. 2016;29: 711–722.

12. Gong J, Mei S, Liu C, Xiang Y, Ye Y, Zhang Z, et al. PancanQTL: systematic identification of cis-eQTLs and trans-eQTLs in 33 cancer types. Nucleic Acids Res. 2018;46: D971–D976.

13. Behrens G, Winkler TW, Gorski M, Leitzmann MF, Heid IM. To stratify or not to stratify: power considerations for population-based genome-wide association studies of quantitative traits. Genet Epidemiol. 2011;35: 867–879.

14. Dimas AS, Nica AC, Montgomery SB, Stranger BE, Raj T, Buil A, et al. Sex-biased genetic effects on gene regulation in humans. Genome Res. 2012;22: 2368–2375.

15. Grossman RL, Heath AP, Ferretti V, Varmus HE, Lowy DR, Kibbe WA, et al. Toward a Shared Vision for Cancer Genomic Data. N Engl J Med. 2016;375: 1109–1112.

16. Webster TH, Couse M, Grande BM, Karlins E, Phung TN, Richmond PA, et al. Identifying, understanding, and correcting technical biases on the sex chromosomes in next-generation sequencing data [Internet]. bioRxiv. 2018. p. 346940. doi:10.1101/346940

17. Andrews S. FastQC A Quality Control tool for High Throughput Sequence Data. In: http://www.bioinformatics.babraham.ac.uk/projects/fastqc/ [Internet]. 2010. Available: http://www.bioinformatics.babraham.ac.uk/projects/fastqc/

18. Bolger AM, Lohse M, Usadel B. Trimmomatic: a flexible trimmer for Illumina sequence data. Bioinformatics. 2014;30: 2114–2120.

19. Kim D, Langmead B, Salzberg SL. HISAT: a fast spliced aligner with low memory requirements. Nat Methods. 2015;12: 357–360.

20. Liao Y, Smyth GK, Shi W. featureCounts: an efficient general purpose program for assigning sequence reads to genomic features. Bioinformatics. 2014;30: 923–930.

21. McKenna A, Hanna M, Banks E, Sivachenko A, Cibulskis K, Kernytsky A, et al. The Genome Analysis Toolkit: A MapReduce framework for analyzing next-generation DNA sequencing data. Genome Res. 2010;20: 1297–1303.

22. DePristo MA, Banks E, Poplin R, Garimella KV, Maguire JR, Hartl C, et al. A framework for variation discovery and genotyping using next-generation DNA sequencing data. Nat Genet. 2011;43: 491–498.

23. Van der Auwera GA, Carneiro MO, Hartl C, Poplin R, Del Angel G, Levy-Moonshine A, et al. From FastQ data to high confidence variant calls: the Genome Analysis Toolkit best practices pipeline. Curr Protoc Bioinformatics. 2013;43: 11.10.1–33.

24. Hinrichs AS, Karolchik D, Baertsch R, Barber GP, Bejerano G, Clawson H, et al. The UCSC Genome Browser Database: update 2006. Nucleic Acids Res. 2006;34: D590–8.

25. Robinson MD, McCarthy DJ, Smyth GK. edgeR: a Bioconductor package for differential expression analysis of digital gene expression data. Bioinformatics. 2010;26: 139–140.

26. Law CW, Chen Y, Shi W, Smyth GK. voom: precision weights unlock linear model analysis tools for RNA-seq read counts. Genome Biol. 2014;15: R29.

27. Xia J, Gill EE, Hancock REW. NetworkAnalyst for statistical, visual and network-based meta-analysis of gene expression data. Nat Protoc. 2015;10: 823–844.

28. Leek JT, Storey JD. Capturing Heterogeneity in Gene Expression Studies by Surrogate Variable Analysis. PLoS Genet. 2007;3: e161.

29. Stegle O, Parts L, Piipari M, Winn J, Durbin R. Using probabilistic estimation of expression residuals (PEER) to obtain increased power and interpretability of gene expression analyses. Nat Protoc. 2012;7: 500–507.

30. Zheng X, Levine D, Shen J, Gogarten SM, Laurie C, Weir BS. A high-performance computing toolset for relatedness and principal component analysis of SNP data. Bioinformatics. 2012;28: 3326–3328.

31. Delaneau O, Ongen H, Brown AA, Fort A, Panousis N, Dermitzakis E. A complete tool set for molecular QTL discovery and analysis. bioRxiv. 2016; doi:10.1101/068635

32. GTEx Consortium. The Genotype-Tissue Expression (GTEx) project. Nat Genet. 2013;45: 580–585.

33. Cavalcante RG, Sartor MA. annotatr: genomic regions in context. Bioinformatics. 2017;33: 2381–2383.

34. Ernst J, Kellis M. ChromHMM: automating chromatin-state discovery and characterization. Nat Methods. 2012;9: 215–216.

35. Kukurba KR, Parsana P, Balliu B, Smith KS, Zappala Z, Knowles DA, et al. Impact of the X Chromosome and sex on regulatory variation. Genome Res. 2016;26: 768–777.

36. Gaykalova DA, Zizkova V, Guo T, Tiscareno I, Wei Y, Vatapalli R, et al. Integrative computational analysis of transcriptional and epigenetic alterations implicates DTX1 as a putative tumor suppressor gene in HNSCC. Oncotarget. 2017;8: 15349–15363.

37. Liu Y, Zheng P. CD24: a genetic checkpoint in T cell homeostasis and autoimmune diseases. Trends Immunol. 2007;28: 315–320.

38. Gou Q, Gong X, Jin J, Shi J, Hou Y. Peroxisome proliferator-activated receptors (PPARs) are potential drug targets for cancer therapy. Oncotarget. 2017;8: 60704–60709.

39. Mayer IA, Arteaga CL. The PI3K/AKT Pathway as a Target for Cancer Treatment. Annu Rev Med. 2016;67: 11–28.

40. Porta R, Borea R, Coelho A, Khan S, Araújo A, Reclusa P, et al. FGFR a promising druggable target in cancer: Molecular biology and new drugs. Crit Rev Oncol Hematol. 2017;113: 256–267.

41. Seshacharyulu P, Ponnusamy MP, Haridas D, Jain M, Ganti AK, Batra SK. Targeting the EGFR signaling pathway in cancer therapy. Expert Opin Ther Targets. 2012;16: 15–31.

42. Demir IE, Tieftrunk E, Schorn S, Friess H, Ceyhan GO. Nerve growth factor & TrkA as novel therapeutic targets in cancer. Biochim Biophys Acta. 2016;1866: 37–50.

43. Zhang Y-L, Wang R-C, Cheng K, Ring BZ, Su L. Roles of Rap1 signaling in tumor cell migration and invasion. Cancer Biol Med. 2017;14: 90–99.

44. Choudhry H, Helmi N, Abdulaal WH, Zeyadi M, Zamzami MA, Wu W, et al. Prospects of IL-2 in Cancer Immunotherapy. Biomed Res Int. 2018;2018: 9056173.

45. Pinho SS, Reis CA. Glycosylation in cancer: mechanisms and clinical implications. Nat Rev Cancer. 2015;15: 540–555.

46. Stowell SR, Ju T, Cummings RD. Protein glycosylation in cancer. Annu Rev Pathol. 2015;10: 473–510.

47. Mehta A, Herrera H, Block T. Chapter Seven - Glycosylation and Liver Cancer. In: Drake RR, Ball LE, editors. Advances in Cancer Research. Academic Press; 2015. pp. 257–279.

48. Liang K-H, Yeh C-T. O-glycosylation in liver cancer: Clinical associations and potential mechanisms. Liver Research. 2017;1: 193–196.

49. Villegas SN, Gombos R, García-López L, Gutiérrez-Pérez I, García-Castillo J, Vallejo DM, et al. PI3K/Akt Cooperates with Oncogenic Notch by Inducing Nitric Oxide-Dependent Inflammation. Cell Rep. 2018;22: 2541–2549.

50. Wang G-L, Iakova P, Wilde M, Awad S, Timchenko NA. Liver tumors escape negative control of proliferation via PI3K/Akt-mediated block of C/EBP alpha growth inhibitory activity. Genes Dev. 2004;18: 912–925.

51. Huang X, Liu G, Guo J, Su Z. The PI3K/AKT pathway in obesity and type 2 diabetes. Int J Biol Sci. 2018;14: 1483–1496.

52. Baar EL, Carbajal KA, Ong IM, Lamming DW. Sex- and tissue-specific changes in mTOR signaling with age in C57BL/6J mice. Aging Cell. 2016;15: 155–166.

