## Supplementary Information for "Distinct molecular etiologies of male and female hepatocellular carcinoma"

### Supplementary Figures

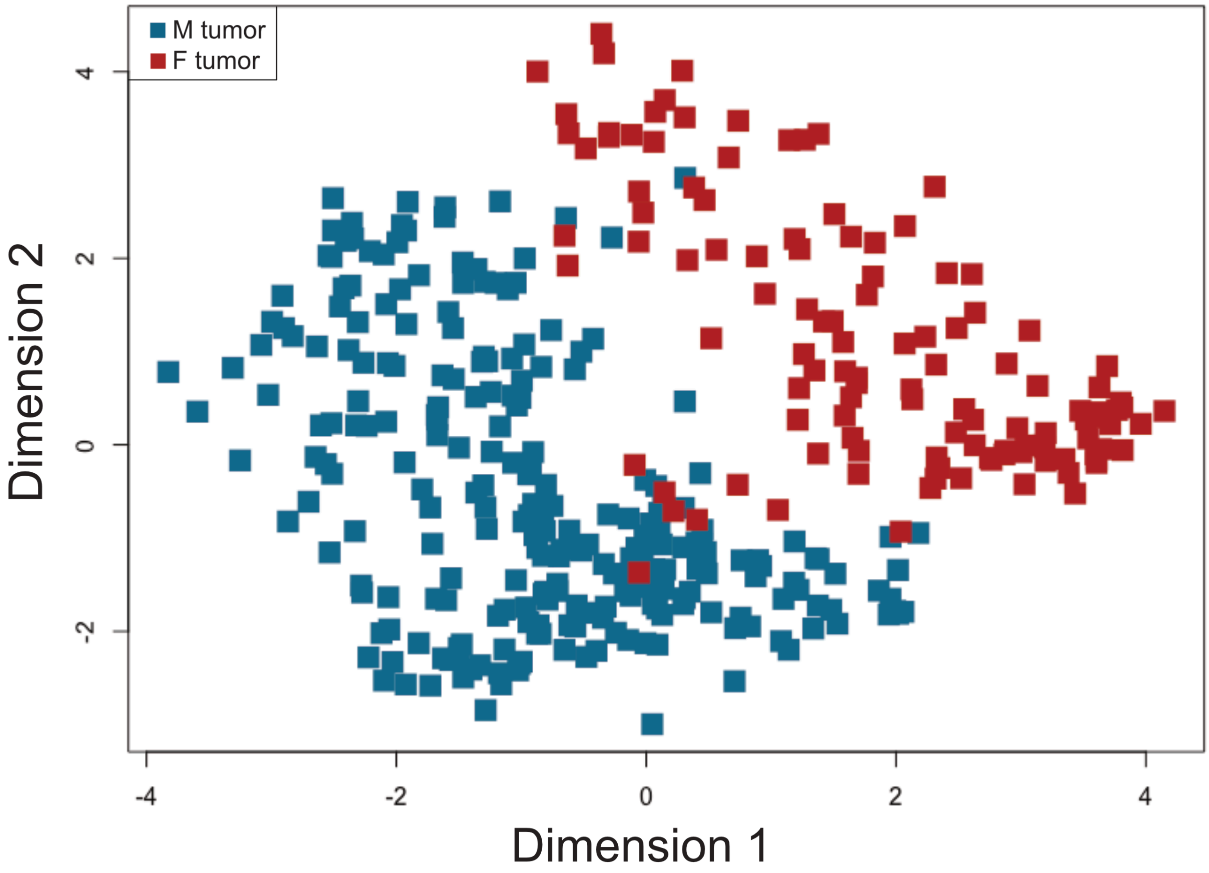


**Fig S1.** A multi-dimensional scaling plot of the TCGA LIHC tumor samples of each sex (N male = 248, N female = 119). Euclidean distances between samples were calculated based on 100 genes with the largest standard deviations between samples.

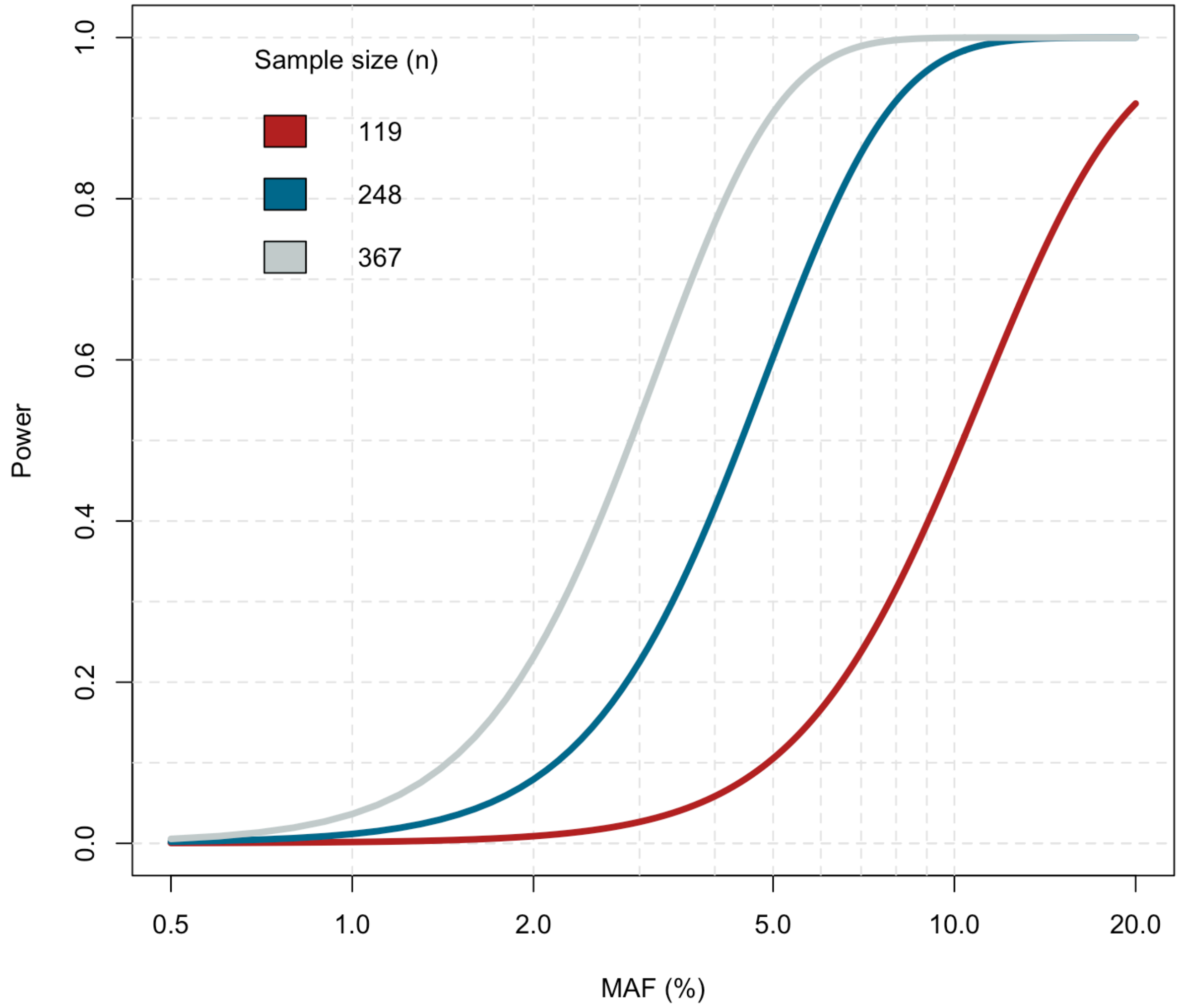


**Fig S2.** Estimation of statistical power in the combined (grey), male-specific (blue), and female-specific (red) eQTL analyses with a *p*-value level 0.01 and 384 variants. Increased power in the combined analysis allows the detection of sex-shared low-effect eQTLs.
